## Supplementary Figures and Table Legends for "Multi-omic analysis of CIC’s functional networks reveals novel interaction partners and a potential role in mitotic fidelity"

### Supplemental Figures

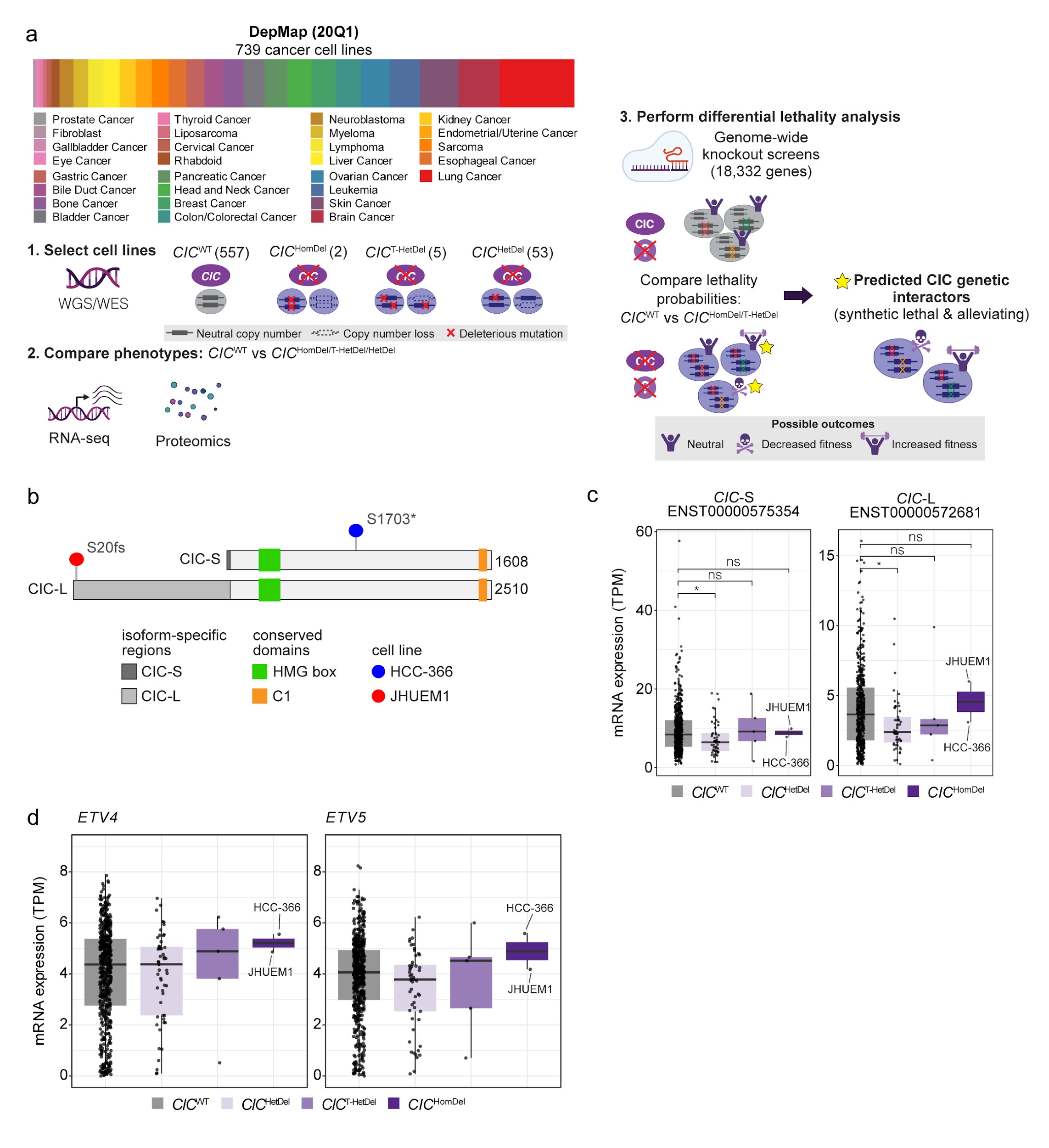

##### **Supplemental Figure S1: Workflow of *CIC* *in silico* genetic interaction screen leveraging data generated by DepMap.**

**a**. Workflow schematic. Cancer cell lines harbouring WT *CIC* alleles (*CIC*^WT^) and mutant *CIC* alleles (*CIC*^HomDel^, *CIC*^T-HetDel^, and *CIC*^HetDel^) were identified using mutation and copy number data from DepMap. mRNA and protein expression data were used to compare *CIC* mutant cell line phenotypes to *CIC*^WT^ phenotypes. The lethality probabilities across 18,333 genes were used to perform Mann-Whitney U-based differential lethality analyses to identify genes with significantly higher or lower lethality probabilities in the *CIC* mutant cell lines compared to *CIC*^WT^ cell lines, which indicate SL or alleviating genetic interactions with *CIC*, respectively. **b.** *CIC* gene model showing the homozygous deleterious alterations identified in both *CIC*^HomDel^ cell lines. **c-d.** *CIC*-S and *CIC*-L isoform transcripts and *ETV4* and *ETV5* total mRNA abundance in *CIC*^HomDel^ (n = 2), *CIC*^T-HetDel^ (n = 5), *CIC*^HetDel^ (n = 53), and *CIC*^WT^ (n =557) cancer cell lines. ANOVA followed by Tukey’s HSD analysis. **p*-value < 0.05 and ns > 0.05.

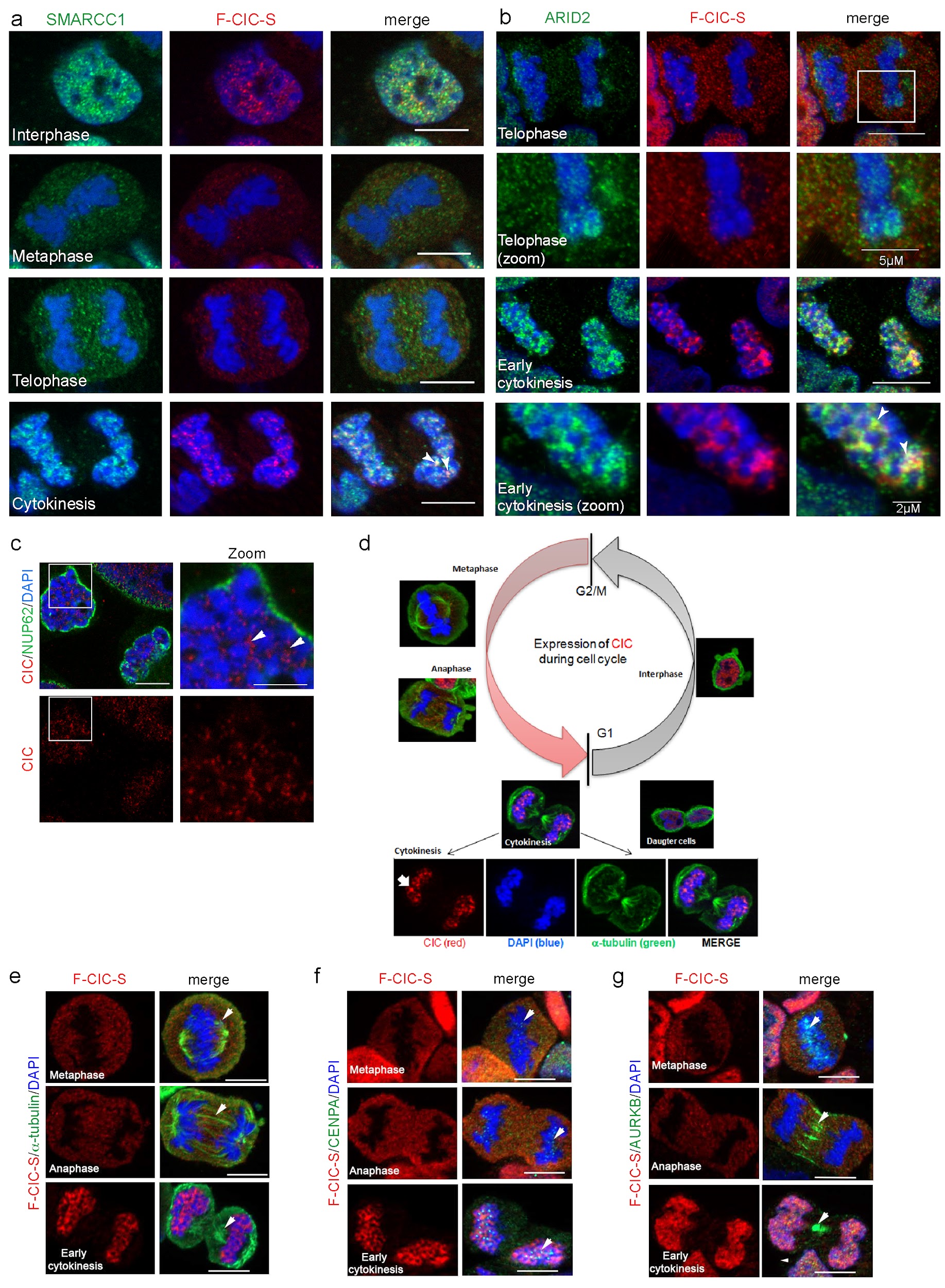

##### **Supplemental Figure S2: Nuclear CIC colocalizes with members of the SWI/SNF complex and is dynamically re-distributed during the cell cycle.**

**a-b**. IF images show colocalization of F-CIC-S (FLAG, red) and SMARCC1 (a) or ARID2 (b, both green) in HEK-*CIC*^KO2 + F-CIC-S^ cells at indicated phases of the cell cycle. DNA was stained with DAPI (blue). Arrowheads indicate colocalization of CIC and the relevant interactor at early cytokinesis (yellow foci). Scale bars: 10 µm or as indicated for zoomed images (b). **c.** Expression of endogenous CIC (red) and the nuclear envelope protein NUP62 (green) in HOG-*CIC*^WT^ cells at cytokinesis. DNA was detected using DAPI staining (blue). Zoomed image of early cytokinesis (right) shows a punctate localization pattern for endogenous CIC throughout the decondensing nucleus (arrowheads). Scale bars: 10 µm and 5 µm (zoomed image). **d**. Expression of F-CIC-S (FLAG, red) and ɑ-tubulin (green) in HEK-*CIC*^KO2^ cells stably expressing F-CIC-S during different stages of the cell cycle. Foci of CIC protein can be observed adjacent to decondensing chromosomes at early cytokinesis (arrow, bottom panel). **e-g** IF images showing no colocalization of F-CIC-S (FLAG, red) with spindles (marked by ɑ-tubulin, e), centrosomes/kinetochores (marked by CENPA, f), or the midbody (marked by AURKB, g) in HEK-*CIC*^KO-D10 + F-CIC-S^ cells. Arrowheads indicate the structures of interest (markers in green), and DNA was stained with DAPI (blue). Representative images are shown for cells in metaphase, anaphase, and early cytokinesis. Scale bar: 10 µm.

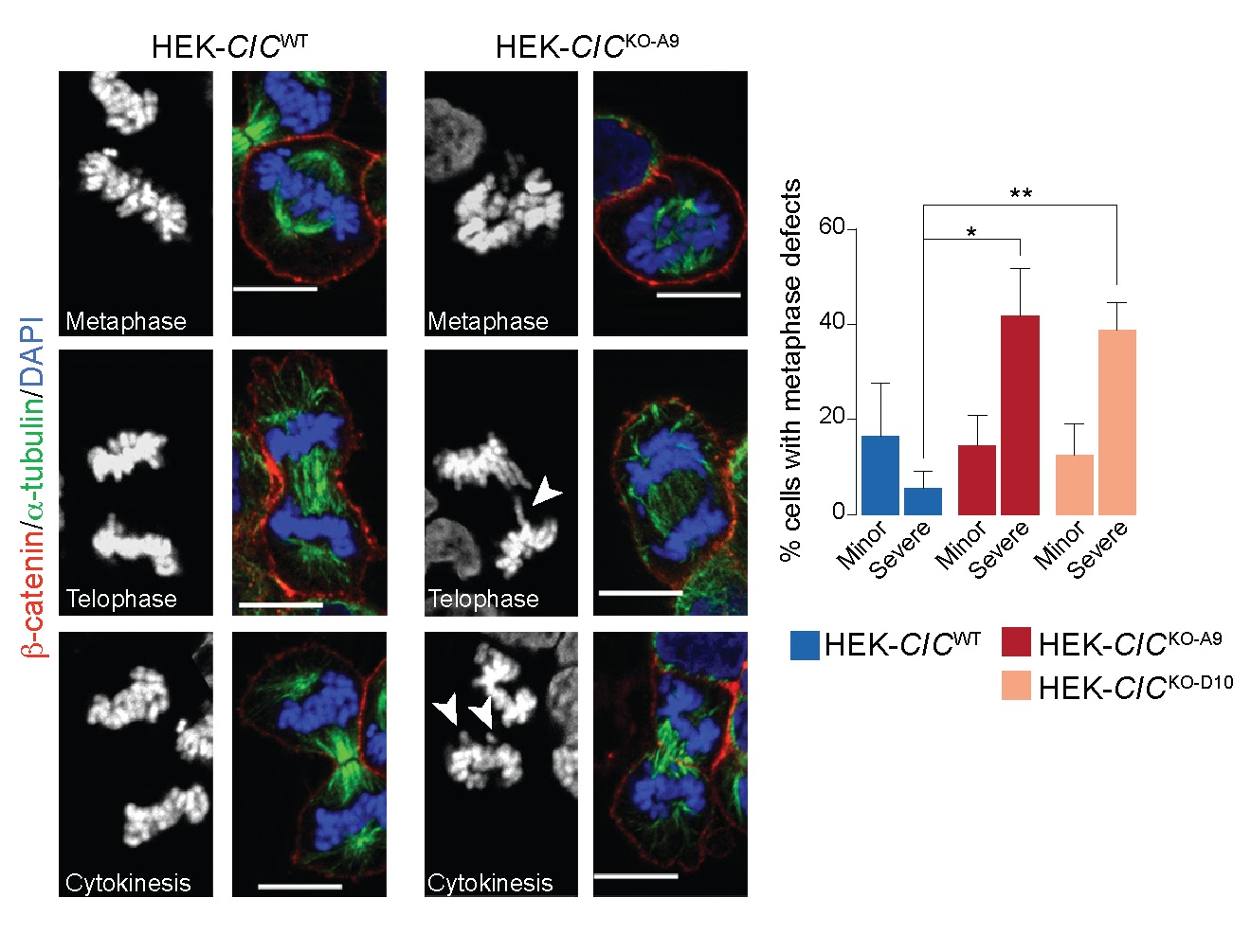

##### **Supplemental Figure S3: Loss of CIC is associated with an increased frequency of mitotic defects.**

Left: representative IF images from HEK-*CIC*^WT^ and HEK-*CIC*^KO-A9^ cells at metaphase and telophase/cytokinesis. Microtubules (ɑ-tubulin, green), DNA (DAPI, blue) and the cell membrane (ß-catenin, red) were detected. DAPI staining alone is shown on the left of each image. HEK-*CIC*^KO-A9^ cells show defects in metaphase alignment and lagging chromosomes at telophase and cytokinesis (arrowheads). Scale bar: 10 µm. Right: bar graphs show proportions of cells with defects at metaphase. Bars represent the mean of three independent experiments and error bars indicate s.e.m. HEK-*CIC*^WT^, n = 59; HEK-*CIC*^KO-A9^, n = 61; HEK-*CIC*^KO-D10^, n = 56. **p*-value < 0.05, ** < 0.01 (two-sided Student’s *t*-test).

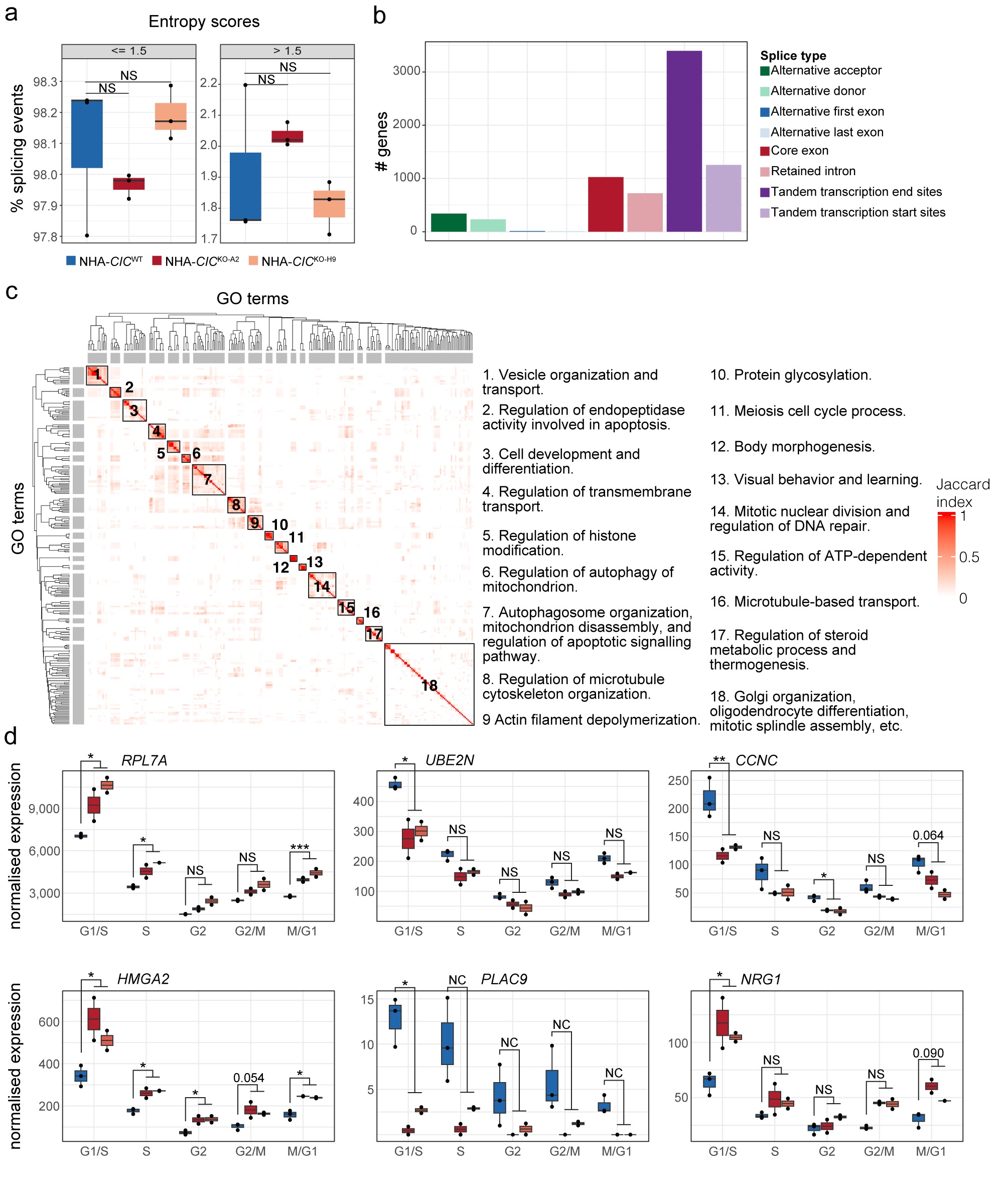

##### **Supplemental Figure S4: Profile of alternative splicing events detected in NHA cell lines.**

**a.** Boxplots showing the percentage of total splicing events detected with either low (<= 1.5; left panel) or high (> 1.5; right panel) entropy scores. Each point represents one of three replicates sequenced from each NHA cell line. NS indicates ANOVA followed by Tukey’s tests that were not significant (p-value > 0.05). **b**. The total number of genes in which differential splicing events were detected between NHA-*CIC*^WT^ and NHA-*CIC*^KO^ cell lines. The splicing events counted includes both significant and non-significant events categorised by type of splicing event. **c**. Heatmap showing Jaccard index similarities between 237 GO terms associated with genes containing significant differential splice events (left). The Jaccard index-based hierarchical clustering was used to summarise the GO terms into 18 groups based on similarity of gene sets associated to the terms, and these functions were summarised in the text (right). **d.** Expression of genes with differential splicing events in RNA-seq data that were also found to be differentially expressed in at least one cell cycle phase in scRNA-seq data. Normalised expression of indicated genes (scRNA-seq data) across cell lines and phases. *BH-adjusted *p*-value < 0.1, ** < 0.01, *** < 0.001 (Wald test as implemented by DESeq2), NS: non-significant, NC: adjusted *p*-value not calculated due to low expression (Methods).

### Supplemental Tables

**Supplemental Table S1: *CIC*’s co-essential and co-expressed gene partners. a** Pearson correlation coefficients were calculated between fitness scores of *CIC* and each targeted gene from DepMap KO

screens. *CIC* co-essential genes were considered as those with a coefficient above the inflection point (0.178) and a BH-adjusted *p*-value < 0.05 (see Methods section for more detail). **b** Co-expression z-scores derived from COXPRESdb’s human Illumina-based RNA-seq data sets.

**Supplemental Table S2: *CIC* mutation characteristics in DepMap 20Q1 cancer cell lines. a-b** Annotations from the DepMap 20Q1 mutations dataset (CCLE_mutations.csv file; **a**) and copy number dataset (CCLE_gene_cn.csv; **b**) for all *CIC* mutations identified across cancer cell lines. Details about the annotations can be found in Ghandi et al. [[70]](https://paperpile.com/c/ussc9G/y7dyL/?noauthor=1). The “CIC isoform” column indicates whether a mutation is in a region specific to either the CIC-L or CIC-S isoforms, or a region common to both. **c** *CIC* mutant group status of all DepMap 20Q1 cancer cell lines (see Methods section for mutant group selection criteria).

**Supplemental Table S3: *In silico* genome-wide screening result comparing *CIC*^HomDel^ mutant cell lines to *CIC*^WT^ control cell lines.** **a** Results obtained from lethality probability comparisons between *CIC*^HomDel^ and *CIC*^WT^ lines. *P*-values were obtained using Mann-Whitney U tests. Adjusted p-values were calculated using permutation tests with 10,000 random samplings. **b** Candidate *CIC* genetic interactors, defined as genes with significant differential lethality probabilities (Adjusted Mann-Whitney U test *p*-value < 0.05 and median lethality probability > 0.5 in at least one group; Methods). **c** GO term annotations for candidate *CIC* genetic interactors shown in b (ClusterProfiler output) and cluster assignment that groups similar terms together (see Supplemental Methods).

**Supplemental Table S4: Genes and proteins identified in the distinct multi-omic analyses in this study. a** Summary table indicating whether a gene or its encoded protein was detected by the corresponding -omics assay indicated by the column name. *CIC* coessential gene: *CIC*’s predicted co-essential gene partners (Supplemental Table S1). *CIC* Genetic Interactor: *CIC*’s synthetic lethal genes predicted in DepMap CIC^HomDel^ mutant cells (Supplemental Table S2). HEK IP-MS: proteins detected in CIC IP-MS data from HEK-*CIC*^WT^ cells (Supplemental Table S5). NHA IP-MS: proteins detected in CIC IP-MS data from NHA-*CIC*^WT^ cells (Supplemental Table S6). scRNAseq differentially expressed genes: Genes that were identified as differentially expressed in at least one cell cycle phase in scRNA-seq analysis comparing NHA-*CIC*^KO^ versus NHA-*CIC*^WT^ cells (Supplemental Table S7). NHA CIC ChIP-seq: high-confidence CIC ChIP peaks identified in Lee et al. [[24]](https://paperpile.com/c/ussc9G/3SqZl/?noauthor=1) (Supplemental Table S9). mESC CIC ChIP-seq: CIC ChIP peaks identified in mESCs profiled by Weissmann et al. [[22]](https://paperpile.com/c/ussc9G/t0Hmc/?noauthor=1) (Supplemental Table S10). Alternatively spliced: genes identified as having differential splicing patterns in NHA-*CIC*^KO^ versus NHA-*CIC*^WT^ cells (Supplemental Table S11). Total # of assays detecting gene: Tally of the number of analyses in which a gene or its encoded protein was detected.

**Supplemental Table S5: Candidate nuclear interactors of CIC in HEK-*CIC*^WT^ cells. a** IP-MS results for candidate CIC interactors identified in at least two of four replicate experiments performed in HEK-*CIC*^WT^ cells (ordered alphabetically). Mascot score: ion score for an MS/MS match based on the calculated probability (see Methods). Total peptides: total number of peptides observed in each IP-MS experiment. Discrete peptides: number of unique peptides observed in each IP-MS experiment. IP: replicate experiment from which data are shown (E1-3: endogenous CIC replicates 1-3, M-1: ectopic MYC-tagged CIC; see Methods). **b** Enrichment analysis results for candidate interactors shown in a (ClusterProfiler output) with cluster assignment grouping similar terms together (see Supplemental Methods).

**Supplemental Table S6: Candidate nuclear interactors of CIC in NHA-*CIC*^WT^ cells. a** Trigger peptides synthesised for NHA-*CIC*^WT^ IP-MS experiment. **b** IP-MS results for NHA-*CIC*^WT^ cells. Proteins detected with a log_2_FC > 0 were considered candidate CIC interactors. PSMs: number of total peptides observed. PEPs: number of unique peptides observed. IgG1-3: protein signal intensity of input control replicates. IP1-3: protein signal intensity of endogenous CIC IP replicates. log_2_FC: log_2_-transformed IP/IgG ratio. **c** High-confidence candidate CIC interactors that were identified in both HEK-*CIC*^WT^ cells (Supplemental Table S4a) and in NHA-*CIC*^WT^ cells (b). **d** Enrichment analysis results for high-confidence candidate interactors shown in c (ClusterProfiler output) with cluster assignment grouping similar terms together (see Supplemental Methods).

**Supplemental Table S7: scRNA-seq phase-specific DE results.** Results (DESeq2 outputs) are shown for NHA-*CIC*^KO^ versus NHA-*CIC*^WT^ comparisons. Base mean: mean of normalised counts for all samples. SE: standard error. Stat: Wald statistic. Adjusted *p*-values were obtained using the Benjamini-Hochberg method. Genes without adjusted *p*-values were filtered by automatic independent filtering for having a low mean normalised count (see Methods).

**Supplemental Table S8: scRNA-seq phase-specific GSEA results.** Results (clusterProfiler outputs) are shown for GSEAs performed on genes ranked according to their differential expression between NHA-*CIC*^KO^ and NHA-*CIC*^WT^ cells.

**Supplemental Table S9: High-confidence CIC ChIP peaks identified in Lee et al.** [[24]](https://paperpile.com/c/ussc9G/3SqZl/?noauthor=1)**.** Genomic coordinates refer to the GRCh37 (hg19) assembly. Peaks were identified using MACS2. Fold-enrichment: read count fold-change between CIC ChIP and matched input. Peaks were annotated using ChIP-seeker, and the nearest gene is indicated. Peaks that overlap with peaks identified by Weissmann et al. [[22]](https://paperpile.com/c/ussc9G/t0Hmc/?noauthor=1) (≥ 1bp) are also indicated.

**Supplemental Table S10: CIC ChIP peaks identified in mESCs.** Genomic coordinates refer to the GRCh37 (hg19) assembly. Peaks were identified using MACS2 using data from Weissmann et al. [[22]](https://paperpile.com/c/ussc9G/t0Hmc/?noauthor=1). Fold-enrichment: read count fold-change between CIC ChIP and matched IgG input. Peaks were annotated using ChIP-seeker, and the nearest gene is indicated. Overlaps with other SWI/SNF peaks (identified using data from Gatchalian et al. [[105]](https://paperpile.com/c/ussc9G/dOfY/?noauthor=1)) are indicated. HGNC symbols are indicated for genes that have human orthologues, as is whether the gene was found to have a nearby CIC ChIP-seq peak in NHA cells [[24]](https://paperpile.com/c/ussc9G/3SqZl).

**Supplemental Table S11: Alternative splicing events detected in NHA cell lines.** **a** Results of all splicing events quantified (Whippet.quant.jl output) in NHA-*CIC*^WT^ line, NHA-*CIC*^KO-A2^ line, and NHA-*CIC*^KO-H9^ line. **b** Results (Whippet.diff.jl output) of differential splicing events detected between NHA-*CIC*^WT^ line and NHA-*CIC*^KO^ lines. Significant splicing events events were those with absolute Delta Ψ > 0.1 and probability > 0.9. **c** GO terms associated with genes affected by significant splicing events (b) were annotated using CluterProfiler. Those with p-values < 0.05 are shown. **d** scRNA-seq phase-specific DE results for the 51 genes affected by significant differential splicing events.

**Supplemental Table S12: Phase-specific genes identified by Whitfield et al.** [[87]](https://paperpile.com/c/ussc9G/olbh5/?noauthor=1) **used to score and assign cells to cell cycle phases.** Gene names from the original study are shown, as well as updated gene names and Ensembl gene IDs used for our data.

**Supplemental Table S13: Antibodies used in this study.**
