## Supplementary Methods and Materials for "Multi-omic analysis of CIC’s functional networks reveals novel interaction partners and a potential role in mitotic fidelity"

### Supplemental Methods

#### Single-cell RNA-seq (scRNA-seq) library construction and sequencing

Cells were harvested using 0.05% Trypsin-EDTA, washed twice with 1x PBS + 0.04% BSA, and passed through a 35 µm strainer to generate a single-cell suspension. Single cell 3' RNA-seq libraries were generated using the Chromium Single Cell 3′ Library & Gel Bead Kit v1 (10X Genomics, Pleasanton, CA, USA) following the manufacturer's protocol. Briefly, the volume of cell suspension was determined based on a targeted cell recovery of 1,000 cells and then mixed with the Single Cell Master Mix prior to loading onto the Single Cell 3' Chip. Following gel bead in emulsion (GEM) generation in the Chromium Controller (10X Genomics), the GEM-RT reaction was performed, GEMs were broken, and cDNA clean up was performed using Dynabeads^TM^ MyOne Silane beads (Thermo Fisher Scientific, Waltham, MA, USA) followed by a cleanup with SPRIselect beads (Beckman Coulter, Indianapolis, IN, USA). cDNA amplification was performed using a total of 14 amplification cycles, followed by reaction clean up with SPRIselect beads. Total cDNA quantification was performed using the Agilent High Sensitivity DNA Kit (Santa Clara, CA, USA). cDNA shearing on the Covaris E220 Focused Ultrasonicator (Woburn, MA, USA), library construction (10 cycles for sample index PCR), and SPRIselect cleanups were performed to generate the final indexed libraries. Library quality was assessed using the Agilent High Sensitivity DNA Kit and quantified using the Qubit dsDNA HS Assay Kit (Thermo Fisher Scientific). Libraries were pooled (8 libraries/pool) and sequenced on one flowcell on the Illumina NextSeq500 platform (San Diego, CA, USA) using paired-end sequencing with dual indexing following Illumina protocols and 10X sequencing run parameters (98bp read1, 14bp i7 index, 8bp i5 index, and 98bp read2).

#### Cell lysis preparations

Cell pellets were resuspended in 4x packed cell volume of RIPA lysis buffer freshly supplemented with 1% sodium orthovanadate (100mM), 1% PMSF (200mM), 2% (v/v) protease inhibitor (sc-24948; Santa Cruz Biotechnology, Dallas, TX, USA), and PhosSTOP (Roche, following the manufacturer’s recommendations). Cell pellets were sonicated briefly and mixed for 1 h at 4°C on a rotator. Cellular debris was removed using centrifugation at 13,000 rpm for 10 min at 4°C. Total protein was quantified using the Pierce™ BCA Protein Assay Kit (Thermo Fisher Scientific).

#### Western blots

Samples were subjected to gel electrophoresis on NuPage^®^ 3-8% Tris Acetate or 10% Bis-Tris pre-cast mini-gels with 1x NuPage^®^ MOPS buffer (Thermo Fisher Scientific). Separated proteins were transferred onto a methanol-activated PVDF membrane (162-0177; Bio-Rad Laboratories, Hercules, CA, USA) in 1x NuPage^®^ Transfer buffer (Thermo Fisher Scientific) + 20% (v/v) methanol. Membranes were blocked with either 2% or 5% (w/v) skim milk in PBST or ReliaBLOT BLOCK (WB 120; Bethyl Laboratories, Montgomery, TX, USA) in TBST for 1 h at room temperature prior to incubation with primary antibodies (Supplemental Table S10) at 4°C overnight. For protein detection, membranes were incubated with HRP-IgG goat α-mouse or α-rabbit (1:5000; Santa Cruz Biotechnology), or ReliaBLOT α-Rabbit HRP (1:5000; Bethyl Laboratories) for 1 h at room temperature followed by three PBST washes before application of ECL substrate (Bio-Rad Laboratories) or SuperSignal West Femto substrate (Thermo Fisher Scientific). Images were captured using a ChemiDoc™ MP Imager and processed with Protein Image Lab 5.1 (Bio-Rad Laboratories).

#### Nuclear fractionation and immunoprecipitation

Cells were grown for 48 h and harvested at ~80% confluency. Fresh cell pellets were gently resuspended in 5x packed cell volume of cytoplasmic lysis buffer (10mM Tris-HCl pH7.2, 10mM NaCl, 2mM MgCl_2_, 1mM EDTA, 0.05% NP-40, with 1X cOmplete protease inhibitor [Roche] and 1X PhosSTOP [Roche]). Lysates were incubated on ice for 10 min before centrifugation at 300g for 5 min to collect nuclear pellets. After 3 to 4 washes with wash buffer (10mM Tris-HCl pH7.2, 10mM NaCl, 2mM MgCl_2_, 1mM EDTA with 1X cOmplete protease inhibitor and 1X PhosSTOP), nuclear pellets were resuspended in 3x packed cell volume of nuclear lysis buffer (250mM NaCl, 20mM sodium phosphate pH7.0, 30mM sodium pyrophosphate, 5mM EDTA, 10mM sodium fluoride, 10% glycerol, 1% NP-40, 1mM DTT, with 1X cOmplete protease inhibitor and 1X PhosSTOP) and then homogenized using a 21-gauge needle. Cellular debris was removed by centrifugation at 13,000 rpm for 30 min at 4°C and then nuclear protein was quantified using the Pierce™ BCA Protein Assay Kit (Thermo Fisher Scientific). Before proceeding with immunoprecipitation, nuclear lysates were treated with 0.1X Benzonase (MilliporeSigma, Burlington, MA, USA) for 30 min at 4°C. Antibody-bound magnetic beads were prepared by incubating anti-CIC or normal rabbit-IgG antibodies (Supplemental Table S10) with Protein G Dynabeads^TM^ (Thermo Fisher Scientific) in PBST (0.1% v/v) at 4°C for 30 min, and then rinsed three times with wash buffer (1X PBS, 1mM EDTA, 0.5% NP40, with 1X cOmplete protease inhibitor and 1X PhosSTOP). Nuclear lysates were incubated with 1.5 mg of the prepared anti-CIC or rabbit IgG (control) Protein-G beads at 4ºC overnight. Protein and protein complexes were released by boiling the magnetic beads in the elution buffer (2X Nupage LDS buffer [Thermo Fisher Scientific], 200mM DTT) at 98^o^C for 10 min.

#### Reciprocal immunoprecipitation

Whole-cell extracts were prepared by lysing fresh cell pellets with 2x packed cell volume of lysis buffer (25mM Tris-HCl pH7.4, 150mM NaCl, 1% NP-40, 5% glycerol, with 1X cOmplete protease inhibitor [Roche] and 1X PhosSTOP [Roche]), followed by homogenization using 10 passes through a 21-gauge needle. After a 30 min incubation on ice, lysis preparations were cleared of cellular debris by centrifugation at 13,000 rpm for 30 min at 4°C and resuspended in 200 ul of lysis buffer. Whole-cell lysates were incubated with anti-ARID1A, anti-ARID2, anti-SMARCA2, anti-SMARCC2, or mouse/rabbit-IgG antibodies (Supplemental Table S10) at 4ºC overnight. Proteins and complexes were captured by incubating antibody-lysate mixtures with Protein G Dynabeads^TM^ for 1 h at 4°C. For elution, magnetic beads were boiled in elution buffer (2X Nupage LDS buffer, 200mM DTT) at 98^o^C for 10 min.

#### HEK cell lines MS data analysis

Data from the 4000 QTrap were processed using Mascot Software (v2.5.1) [[155]](https://paperpile.com/c/ussc9G/xr9K). MS2 spectra were searched against the Uniprot-Swissprot database (v2020March) [[156,157]](https://paperpile.com/c/ussc9G/NX2q+VFWc) using the Homo sapiens taxonomy filter (20,366 total entries). Mascot parameters were specified as: trypsin enzyme, 1 missed cleavage allowed, peptide mass tolerance of 0.8 Da, and a fragment mass tolerance of 0.8 Da. Oxidation of methionine and deamidation at NQ were set as variable modifications. Carbamidomethylation of cysteine was set as a fixed modification. An ion score cutoff of 34 and required bold red criteria were used to filter the protein hits.

#### NHA cell lines LC-MS/MS analysis

##### Protein elution, clean-up with SP3, and protease digestion

Proteins eluted from IP preparations from NHA lines in SDS loading buffer were purified using the SP3 method, as described previously [[158,159]](https://paperpile.com/c/ussc9G/Tkaf4+rmDsO). A 1:1 combination of two different types of carboxylate-functionalized beads, both with a hydrophilic surface (Sera-Mag Speed Beads, 45152105050350 and 65152105050350; GE Life Sciences, Chicago, IL, USA) was added to the lysate and ethanol was added to achieve a final concentration of 50% by volume. Tubes were mixed on a ThermoMixer unit (Eppendorf, Hamburg, Germany) at 1,000 rpm for 10 min at room temperature, then placed in a magnetic rack and incubated for 2 min. The supernatant was discarded, and the beads were rinsed 3x with 180 μL of 90% ethanol by removing the tubes from the magnetic rack and gently re-suspending the beads by pipette mixing. For elution, the tubes were removed from the magnetic rack and beads were resuspended in 100 μL of 50 mM HEPES (pH 8) containing an appropriate amount of trypsin/rLysC mix (1:25 enzyme to protein concentration; V5071; Promega, Madison, WI, USA) and incubated for 14 h at 37**°**C in a ThermoMixer with mixing at 1000 rpm. After incubation, the tubes were sonicated briefly (30 s) in a bath sonicator, placed on a magnetic rack, and the supernatant was recovered for further processing.

##### Synthetic peptide mix preparation

The set of standard peptides was taken as a subset from the collection analyzed in the ProteomeTools initiative [[160]](https://paperpile.com/c/ussc9G/ebRsP). Peptides were selected for a panel of 51 proteins, resulting in a set of 254 total candidates that were synthesized in a ‘SpikeMix’ format (Supplemental Table S6; JPT Peptide Technologies, Berlin, Germany). Upon delivery, dried peptides were reconstituted in 100 μL of DMSO, vortexed briefly (~15 s), and sonicated in a water bath for 5 min. Reconstituted peptides were measured in a dilution series to determine the concentration that represented the limit of detection for the majority of the pool. Reconstituted peptides were spiked into interactome samples at a concentration 10% above the determined limit of detection. In this way, the synthetic spikes would not negatively impact the resulting quantification of detected peptides due to a large difference in dynamic range in comparison to the IP samples.

##### Tandem mass tag (TMT) labelling of peptides

TMT 6-plex labelling kits were obtained from Pierce (Thermo Fisher Scientific). Each TMT label (5 mg per vial) was reconstituted in 500 μL of acetonitrile and refrozen. Labelling reactions were carried out through addition of 200 μg of TMT label in two volumetrically equal steps of 10 μL (100 μg per addition), 30 min apart. Control IP samples were labelled with TMT 126C, 127N, and 128C. Target IP samples were labelled with TMT 129N, 130C, and 131N. The synthetic peptide spikes were labelled using the TMT131C reagent (from the TMT 11-plex reagent set). Reactions were quenched by the addition of 10 μL of glycine (1M stock solution, Sigma-Aldrich). Labelled peptides were concentrated on a SpeedVac centrifuge (Thermo Fisher Scientific) to remove excess acetonitrile, acidified to 1% (v/v) trifluoroacetic acid (TFA), and purified with C18 StageTips (Thermo Fisher Scientific).

##### Peptide clean-up procedures

Peptides were desalted and concentrated using StageTip treatment as described previously [[161]](https://paperpile.com/c/ussc9G/9J04m). For StageTip clean-up, three discs of C18 Empore material (66883-U; Sigma-Aldrich) packed in 200μL pipette tips were rinsed twice with 100 μL of acetonitrile with 0.1% TFA. Cartridges were then rinsed twice with 100 μL of water with 0.1% TFA prior to sample loading. Loaded samples were rinsed twice with 0.1% formic acid (100 μL per rinse) and eluted with 100 μL of 80% acetonitrile containing 0.1% formic acid. All TopTip- or StageTip-processed samples were concentrated in a SpeedVac centrifuge (Thermo Fisher Scientific) and subsequently reconstituted in 1% formic acid with 1% DMSO in water.

##### MS analysis of peptide samples on the Orbitrap Fusion

Analysis of TMT-labelled peptide pools was carried out on an Orbitrap Fusion Tribrid MS platform (Thermo Fisher Scientific). Samples were introduced using an Easy-nLC 1000 system (Thermo Fisher Scientific). Columns used for trapping and separations were packed in-house. Trapping columns were packed in 100 μm internal diameter capillaries to a length of 25 mm with C18 beads (Reprosil-Pur, 3 μm particle size; Dr. Maisch HPLC GmbH, Ammerbuch, Germany). Trapping was carried out for a total volume of 10 μL at a pressure of 400 bar. After trapping, gradient elution of peptides was performed on a C18 column (Reprosil-Pur, 1.9μm particle size; Dr. Maisch HPLC GmbH) packed in-house to a length of 20 cm in 100 μm internal diameter capillaries with a laser-pulled electrospray tip and heated to 50**°**C using AgileSLEEVE column ovens (Analytical Sales & Services, Flanders, NJ, USA). Elution was performed with a gradient of mobile phase A (water and 0.1% formic acid) to 8% B (acetonitrile and 0.1% formic acid) over 5 min, to 30% B over 88 min, and to 40% B over 19 min, with final elution (80% B) using a further 8 min at a flow rate of 350 nL/min.

Data acquisition on the Orbitrap Fusion was carried out using a data-dependent method with multi-notch synchronous precursor selection MS3 scanning for TMT tags. Survey scans covering the mass range of 350 – 1500 were acquired at a resolution of 120,000 (at m/z 200), with quadrupole isolation enabled, an S-Lens RF Level of 60%, a maximum fill time of 50 ms, and an automatic gain control (AGC) target value of 5e5. For MS2 scan triggering, monoisotopic precursor selection was enabled, charge state filtering was limited to 2 – 4, an intensity threshold of 5e3 was employed, and dynamic exclusion of previously selected masses was enabled for 60 s with a tolerance of 20 ppm. MS2 scans were acquired in the ion trap in Rapid mode after CID fragmentation with a maximum fill time of 150 ms, quadrupole isolation, an isolation window of 1.6 m/z (0.2 m/z offset), collision energy of 30%, activation Q of 0.25, injection for all available parallelizable time turned OFF, and an AGC target value of 4e3. Fragment ions were selected for MS3 scans based on a precursor selection range of 400-1600 m/z, ion exclusion of 20 m/z low and 5 m/z high, and isobaric tag loss exclusion for TMT. The top 10 precursors were selected for MS3 scans that were acquired in the Orbitrap after HCD fragmentation (NCE 60%) with a maximum fill time of 150 ms, 50,000 resolution, 120-750 m/z scan range, ion injection for all parallelizable time turned OFF, and an AGC target value of 1e5. The total allowable cycle time was set to 4 s. MS1 and MS3 scans were acquired in profile mode, and MS2 in centroid format.

##### NHA cell lines MS data analysis

Data from the Orbitrap Fusion were processed using the Proteome Discoverer Software (v2.1.1.21). MS2 spectra were searched using Sequest HT against a combined UniProt human proteome database appended to a list of common contaminants (24,624 total sequences). Sequest HT parameters were specified as: trypsin enzyme, 2 missed cleavages allowed, minimum peptide length of 6, precursor mass tolerance of 20 ppm, and a fragment mass tolerance of 0.6. Oxidation of methionine and TMT 6-plex at lysine and peptide N-termini were set as variable modifications. Carbamidomethylation of cysteine was set as a fixed modification. Peptide spectral match error rates were determined using the target-decoy strategy coupled to Percolator modeling of positive and false matches [[162,163]](https://paperpile.com/c/ussc9G/29yMd+RKa5x). Data were filtered at the peptide spectral match-level to control for false discoveries using an adjusted *p*-value cut off of 0.01 as determined by Percolator. Contaminant and decoy proteins were removed from all data sets prior to downstream analysis.

#### Immunofluorescence assays

For cell lines, fixed cells were permeabilized using 0.2% TritonX-100/PBS for 5 min, blocked with 2.5% goat serum (G6767; Sigma-Aldrich) in PBS for 1 h on a rocker, and stained with appropriate primary antibodies (Supplemental Table S13) in 0.25% goat serum in PBS overnight at 4°C. Alexa Fluor^®^ 488 and/or 568 (Thermo Fisher Scientific) were used as secondary antibodies (Supplemental Table S13), followed by mounting with DAPI (D3571; Thermo Fisher Scientific) and SlowFade^™^ Diamond Antifade Mountant (36967; Thermo Fisher Scientific).

For mouse tissue, embryonic (E13.5) mouse brain section slides were quickly pre-washed in 1X TBST (0.1% Tween) followed by a 15 min permeabilization in TBST (0.25% Tween) and an additional 5 min wash in TBST (0.1% Tween) with gentle shaking. Slides were blocked with 3% goat serum in 1X PBST (0.1% Tween), covered with parafilm, for 30 min in a humidified staining chamber. α-Phospho Histone H3 (Cell Signaling Technology, Danvers, MA, USA) and α-Rabbit Alexa Fluor^®^ 568 (Thermo Fisher Scientific) antibodies were applied in blocking solution overnight at 4°C and for 1 h at room temperature, respectively, with three 5 min TBST (0.1% Tween) washes performed in between. Slides were then mounted for imaging and quantification. All steps were performed at room temperature unless indicated otherwise.

#### Proximity ligation assays (PLAs)

Cells were plated on an 8-well CC_2_ chamber slide with cover (12-565-1; Fisher Scientific) in growth media. After 19 h, cells were fixed with 4% PFA. PLAs were performed using the Duolink^®^ In Situ Red Starter Kit Mouse/Rabbit (DUO92101; Sigma-Aldrich), following the manufacturer’s protocol. High-resolution images were obtained with an Eclipse Ti inverted confocal microscope equipped with A1 si laser (Nikon, Tokyo, Japan) and Rolera EM-C^2TM^ (Teledyne QImaging, Surrey, BC, Canada) camera. Images used for quantifications were obtained with an Axio Observer inverted fluorescent microscope (Zeiss, Oberkochen, Germany), equipped with Apotome.2 and AxioCam MRm camera. Images were obtained at 40x or 63x (oil) magnification using the Zen software (2.3, blue edition; Zeiss). PLA dots were counted using Blob-Finder software (<http://www.cb.uu.se/~amin/BlobFinder/>).

#### Human chromosomal defect counts

To quantify cells for metaphase defects, 20-25 images per replicate were captured with 40x magnification. Normal, minor defective (a few alignment defects), and moderate-severe defective (severe alignments and multipolar alignments) phenotypes were observed. The proportion of cells with a defective phenotype (number of cells with defective phenotypes in metaphase / total number of cells in metaphase) was calculated for each image. Similarly, for cytokinesis defects, 50 images per replicate were captured with 63x (oil) magnification and the proportion of cells with a defective phenotype (number of cells with defective phenotypes in cytokinesis / total number of cells in cytokinesis) was calculated for each image. We did not observe a sufficient number of cells in cytokinesis in the HEK lines to quantify defects. All images were captured with a Zeiss Axio Observer with Apotome.2 fluorescence microscope.

#### Mouse chromosomal defect counts

E13.5 mouse brain [[84]](https://paperpile.com/c/ussc9G/fyfvZ) section slides were mounted on glass slides and stored at −80°C until analysis. Each slide contained four consecutive sections each from two mice, for a total of eight sections per slide. α-tubulin (Abcam, Cambridge, UK) and DAPI staining were used to quantify chromosomal defects and α-phospho-Histone H3 (Cell Signaling Technology) staining was used to visualize mitotically active cells. For each brain section, images from 10 regions with cells in anaphase and telophase were captured at 63x (oil) magnification using an Axioplan microscope. The proportion of cells with defective anaphase or telophase to the total number of cells at anaphase or telophase, respectively, was used for quantifications.

#### Microscopy imaging

Microscope information is included in relevant sections. Raw images were further processed in Photoshop (Adobe, Mountain View, CA, USA).

#### Enrichment analysis and gene function annotation

We used (v3.16.1) (Wu et al. 2021) to identify pathway and protein complexes significantly enriched (BH-adjusted *p*-value < 0.05) in *CIC* genetic interactors and protein interactors, as well as to annotate functions of gene affected by splicing events with GO biological pathways (unadjusted p-value < 0.05). Given many related GO terms with similar sets of genes associated with them, we calculated the degree of overlapping genes associated between GO terms using the Jaccard index, then summarised them into distinct groups by performing hierarchical clustering of the Jaccard index between pairs of GO terms. The number of distinct clusters was determined using the gap statistic, which calculated the optimal number of clusters (up to 30 clusters) by iteratively bootstrapping 1000 times using the cluster (v2.1.4).
